# Complementary cortical and striatal encoding of locomotor preparation and performance

**DOI:** 10.1101/2025.04.24.650464

**Authors:** Deepak Singla, Long Yang, Andrew S. Weakley, Dylan Davidoff, Jonathan C. Kao, Sotiris C. Masmanidis

## Abstract

Cortical and striatal circuits play an important role in motor planning and execution, and encode movement-related information. While neural dynamics in these areas show substantial similarities, possibly reflecting shared information content, studies directly comparing the cortical and striatal encoding of locomotor preparation and performance have been lacking. Here we contrasted the neural coding properties of mouse primary motor and medial prefrontal cortex, as well as dorsolateral and dorsomedial striatum, prior to and during bouts of self-initiated walking. All four areas contained cells active during both the preparatory and performance periods of locomotion. However, the decoding of behaviorally relevant information using population-level activity revealed significant regional variations. Specifically, dorsomedial striatum more accurately encoded the preparatory period prior to walking, while primary motor cortex more accurately encoded rhythmic limb kinematics during walking. Together, this work provides evidence for a complementary neural coding scheme for locomotor preparation and performance in cortical and striatal circuits.

## Introduction

From flies to mice to humans, walking is a ubiquitous mode of locomotion characterized by a rhythmic pattern of limb movements known as gait. Yet despite the prevalence and simplicity of this behavior relative to other more widely studied motor skills, the neural dynamics underlying locomotion are not fully characterized^1-4^. Fundamentally, the control of all forms of walking involves two critical stages: a preparatory phase that precedes the onset of movement, and a performance phase in which the gait cycle is executed. Our understanding of the brain circuits governing these two stages of locomotion has greatly expanded with the use of neural recordings in animals performing instructed or self-initiated movements. The onset of movement is preceded by preparatory neural activity which emerges hundreds of milliseconds before muscle activation^5-7^. Intriguingly, parallel work has revealed that some of the same cortical and subcortical circuits implicated in movement preparation also exhibit activity related to the continuous production of various movements, including walking^7-10^.

This potential functional overlap is particularly evident from studies of corticostriatal circuit activity. Both cortex and striatum contain neurons displaying preparatory and locomotion-related dynamics^10-19^. And indeed, movement-related activity patterns in these regions appear remarkably similar^20-22^. This similarity may arise via feedforward corticostriatal interactions^23-26^, or via basal ganglia-thalamo-cortical feedback loops^27-30^. In Parkinson’s disease patients and animal models, altered cortical and striatal activity has been linked to impairments in both initiating and performing locomotion^17, 31^. Collectively, these findings suggest that the neural representation of both movement preparation and ongoing locomotion may largely be indistinguishable in cortex and striatum. But crucially, the extent of this similarity has not been directly quantified at the level of neural population dynamics. Thus, it remains unclear whether both circuits contain identical neural codes, or if the strongest representations of locomotor preparation and performance are regionally segregated^32, 33^.

To address this gap, we compared preparatory and performance-related neural dynamics in two cortical and two striatal regions: primary motor cortex (M1), medial prefrontal cortex (mPFC), as well as dorsolateral and dorsomedial striatum (DLS and DMS)—areas implicated in motor planning and execution^11, 12, 34-36^. These circuits are likely to be functionally related due to anatomical connections, with M1 projecting to DLS and mPFC projecting to DMS^23, 24^. Single-unit recordings were carried out in mice performing bouts of spontaneous, self-initiated walking in an open field. We also monitored limb movements using high speed video and automated pose tracking tools. Population decoding methods were then applied to assess how accurately each area represented the time period prior to movement initiation as well as during locomotion. Surprisingly, the most accurate representation of movement preparation was found in DMS. By contrast, information about the locomotor rhythm (i.e., the phase of limbs undergoing gait) was most accurately decoded from M1. Together, these findings have two key implications for understanding the role of cortical and striatal circuits in motor control. First, the results suggest that while activity in these circuits may appear similar superficially, significant differences in their information coding properties are present. Second, this work provides evidence that cortical and striatal circuits serve complementary roles in signaling the preparation and performance of self-initiated locomotion.

## Results

### Distributed representation of locomotion in corticostriatal circuits

Electrophysiological recordings in M1, mPFC (including secondary motor cortex, M2), DLS and DMS were carried out with implanted silicon microprobes as mice freely behaved in an open field, while simultaneously capturing limb movements at high spatial and temporal resolution (**Fig. 1a**). Animals initiated locomotion spontaneously in the absence of sensory triggers or explicit rewards, performing bouts of walking in which individual limbs displayed characteristic gait patterns. To ensure consistency over multiple trials, the onset of each walking bout was aligned to the peak in whole-body acceleration (**Fig. 1b**). Our analysis first investigated the temporal structure of neural spiking around the onset of walking. Across multiple animals and sessions, we recorded a total of at least 200 single-units from each of the four selected brain areas (**Fig. 1c**). Consistent with previous studies in these circuits^12, 13^, we identified a subset of cells displaying ramp-like changes in mean firing rate beginning hundreds of milliseconds before the start of walking, and peaking shortly after movement onset before gradually decreasing (**Fig. 1d,e**). As a preliminary measure of the dynamics that may be related either to movement preparation or performance, we assessed the proportion of task-modulated cells around the time of walking onset^37^. While superficially, similar trends were found in all four regions, some potential variability was observed (**Fig. 1f**). However, it is difficult to draw firm conclusions from these results. For example, it is conceivable for a circuit to contain a sparse but informative representation of locomotor preparation or performance. Furthermore, from a computational standpoint, contemporary work has stressed the importance of population-level dynamics, rather than individual neurons, in the preparation and performance of movement^38, 39^. Thus, while analyzing the average firing properties of individual cells is useful for understanding the overall prevalence of locomotor activity, these results do not necessarily reflect the trial-to-trial variability of information encoded at the population level.

**Fig. 1:**
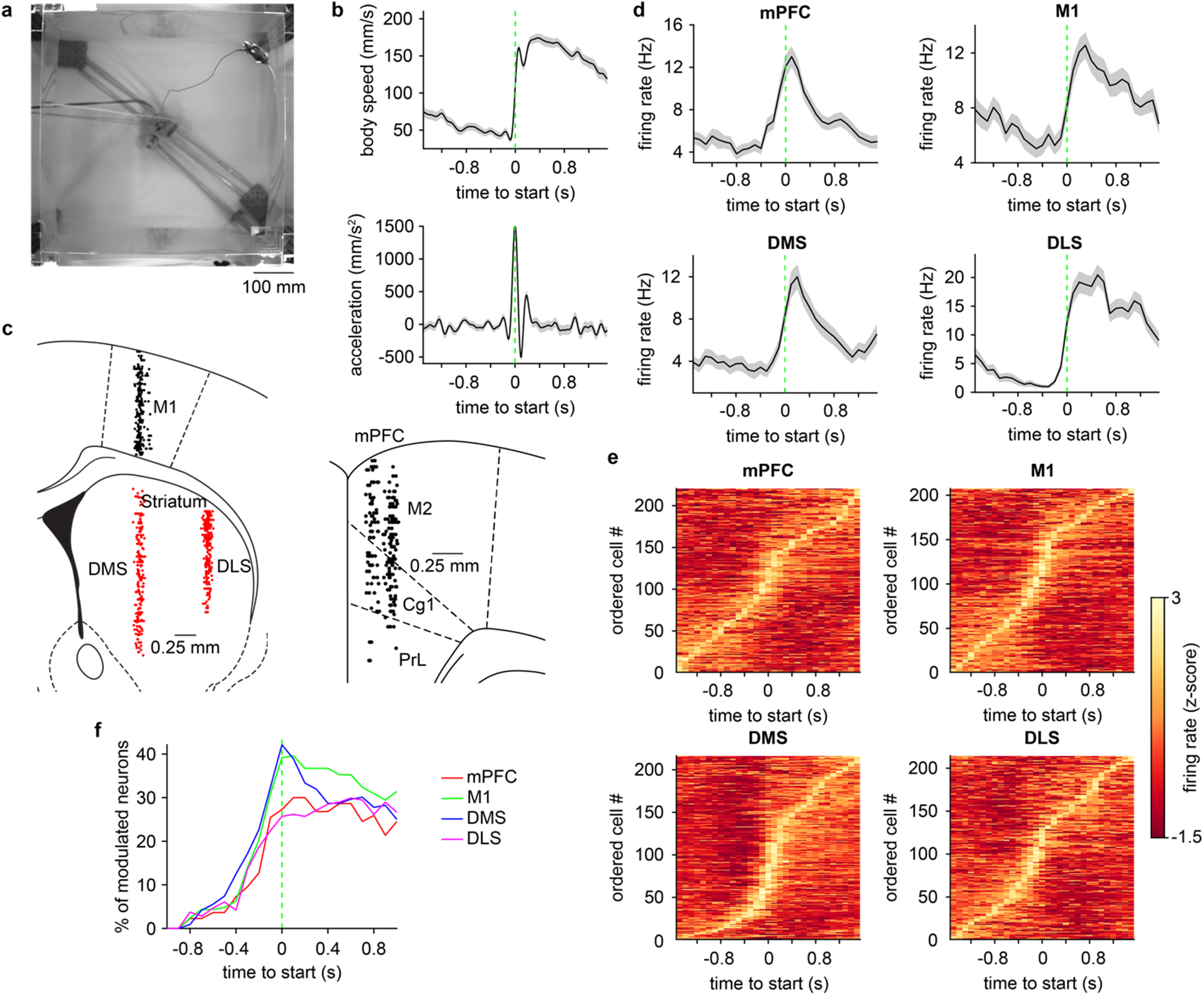
Single unit activity in cortical and striatal circuits during self-initiated locomotion. **a,** A video frame showing the bottom-up view of a mouse in the experimental arena. **b,** Mean body speed (top) and acceleration (bottom) of one animal around the start of walking. Green dashed line represents the onset of walking. **c,** Left, location of single units recorded from M1 (n = 207 cells pooled from 21 sessions in 7 mice) in black, DMS (n = 216 cells pooled from 10 sessions in 4 mice) and DLS (n = 214 cells pooled from 11 sessions in 4 mice) in red. Right, location of single units recorded from mPFC (n = 220 cells pooled from 9 sessions in 2 mice). M2: secondary motor, Cg1: primary cingulate, PrL: prelimbic cortex. **d,** Mean firing rate of a representative cell from each of the four areas aligned to the start of walking. **e,** Mean firing rate of all cells from each area aligned to the start of walking. The cells are ordered by the time of maximum firing. **f,** Percentage of cells modulated in each of the areas aligned to the start of walking. Shading in **b** and **d** represents ± s.e.m.

### More accurate decoding of locomotor preparation from striatal dynamics

To address potential limitations with interpreting single-cell average firing properties, we next examined population-level activity around the start of locomotion. We tested whether a decoder could be trained on activity from each area to correctly identify different time points spanning both the preparatory and performance phases of movement (**Fig. 2a**). This approach enabled us to assess how consistently neural activity at each time point could be distinguished from activity at every other time point on an individual trial basis^40^. In general, population activity close to movement onset could be identified most accurately (**Fig. 2b**). Conversely, activity became less distinguishable at progressively earlier and later times relative to movement onset^37^. The decoders performed better than chance levels for multiple time steps prior to movement onset, suggesting an accurate representation of locomotor preparation (**Fig. 2c**). We also confirmed that the decoding accuracy improved with the number of cells used to train the classifier (**Extended Data Fig. 1a,b**). Importantly, although these trends were qualitatively similar across all four recorded areas, when comparing decoders trained on equivalent numbers of cells, the accuracy levels displayed marked regional variations. We quantified these differences by separately calculating the average decoding accuracy in the pre- and post-start time periods, corresponding to the preparatory and performance phases of walking. In the pre-start (i.e., preparatory, -0.9 to - 0.3 s) period, decoding was significantly more accurate in the DMS population, followed by DLS (**Fig. 2d**). By contrast, in the post-start (i.e., performance, 0.3 to 0.9 s) period when locomotion was underway, decoding from the M1 population significantly outperformed the other three areas.

**Fig. 2:**
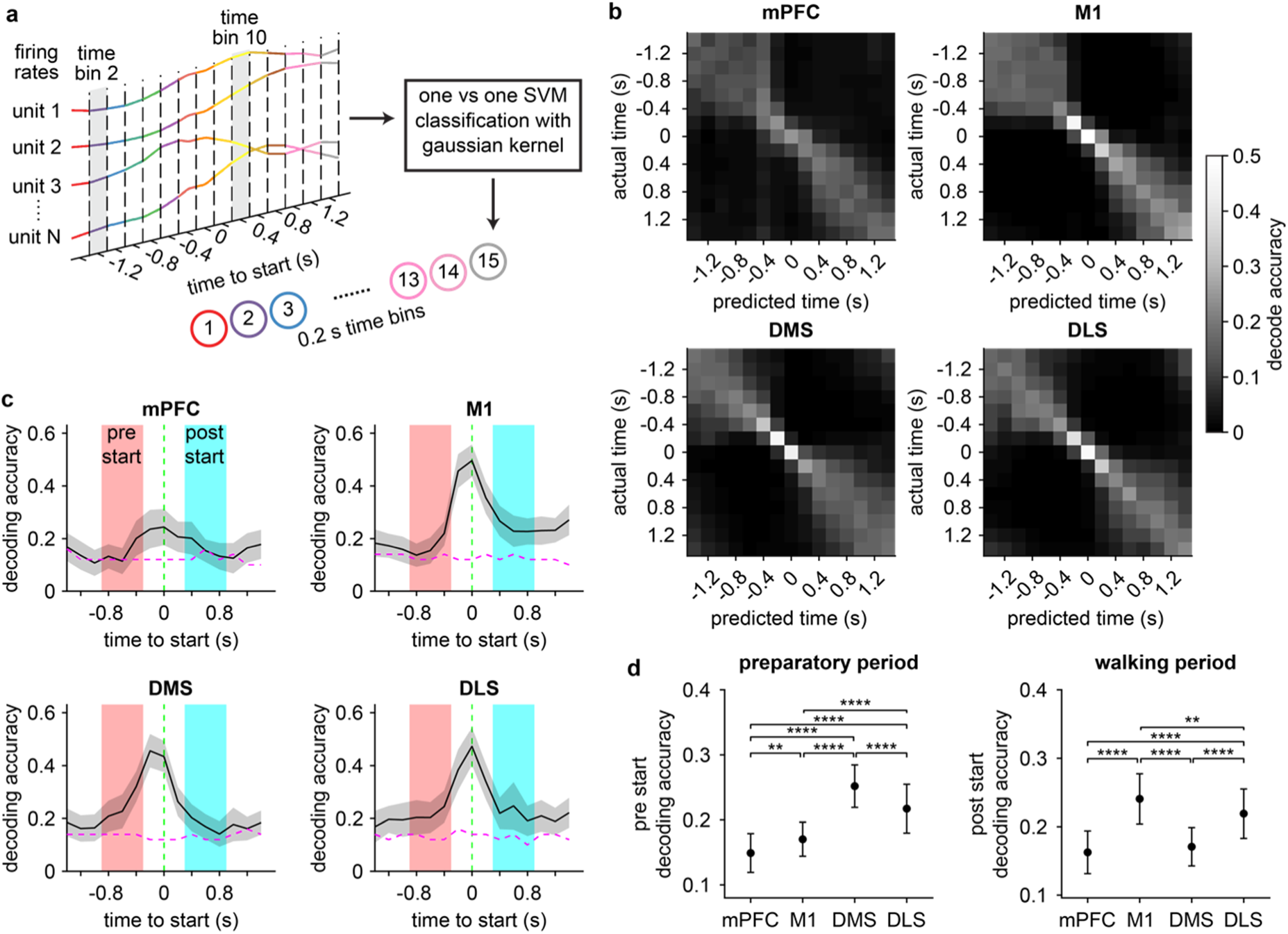
Regional differences in decoding locomotor preparation and performance from population dynamics. **a,** Schematic of the approach used to decode time around the start of walking using a support vector machine (SVM) classifier. Firing rate estimates of single units are binned in 0.2 s increments for a total of 15 population activity patterns per trial. The decoder is trained to distinguish activity patterns in each time bin from every other time bin. **b,** Confusion matrices displaying how well the decoder performed in each of the four areas. **c,** Mean decoding accuracy aligned to the start of walking. Magenta dashed line indicates the 95 % confidence interval of decoder performance trained on time-shuffled data. Green dashed line represents the onset of walking. **d,** Left, mean decoding accuracy in the preparatory (“pre-start”, t = -0.9 to -0.3 s) period across the four areas (one-way ANOVA, n = 50 random drawings of 200 cells, F_3,196_ = 106, p < 0.0001). Right, mean decoding accuracy in the performance (“post-start”, t = 0.3 to 0.9 s) period of walking across the four areas (one-way ANOVA, n = 50 random drawings of 200 cells, F_3,196_ = 64, p < 0.0001). Data in **c** and **d** represents the mean ± s.d. of 50 random drawings of 200 cells. Post hoc Tukey-Kramer test was applied for all area-wise comparisons.

To verify these results, we applied a separate measure of decoder performance, the root mean squared error (RMSE) between the predicted and actual time points (i.e., lower RMSE values reflect better decoding). The RMSE was significantly lower in DMS and DLS during the pre-start period, but significantly lower in M1 during the post-start period (**Extended Data Fig. 1c,d**). Taken together, these findings suggest that striatal population dynamics, particularly in DMS, contain a more accurate temporal code than M1 in the preparatory phase of self-initiated locomotion. However, this relationship appears to be reversed as animals perform walking, with M1 coding surpassing DMS. Temporal coding in DLS was generally of intermediate quality between M1 and DMS. Unexpectedly, mPFC recordings, which included M2, consistently showed the lowest decoding accuracy in the preparatory period despite being frequently implicated in motor planning^35, 36^.

### Stronger population code for whole-body speed in M1 and DLS

The post-start period consists of walking behavior. We therefore sought additional clarity on the finding that temporal coding in this time period was more accurate in M1 and DLS. Since whole-body speed is a widely used, albeit simple, measure of locomotion^14, 41, 42^, we initially examined speed coding during bouts of walking at both the single cell and population level. A subset of cells from each area showed speed-related changes in firing rate (**Fig. 3a**). Since firing rate often appeared to approximately vary linearly with speed, we assessed the strength of this relationship by correlating each cell’s firing rate with speed^42^. This analysis revealed a significantly higher average speed correlation, and a higher proportion of speed correlated cells in DLS compared to mPFC (**Fig. 3b,c**). However, these differences were relatively modest in size, with over 50 % of cells in each area showing significant correlation to speed. It has been previously recognized that this single cell correlation-based analysis approach may not capture the complex relationship between neural population dynamics and body speed^41^. Thus, we again turned to population decoding methods to classify neural activity during walking into one of four speed ranges (50 to 250 mm/s in 50 mm/s increments). For all four areas, speed decoding was moderately successful, with only some speed ranges being predicted above chance levels (**Fig. 3d**). However, on average the speed decoding accuracy was significantly higher in M1 and DLS compared to DMS and mPFC (**Fig. 3e**). Likewise, the decoding error (RMSE) was significantly lower in M1 and DLS (**Fig. 3f**). These results suggest that M1 and DLS contain a more informative population code for whole-body speed, a simple readout of locomotor performance.

**Fig. 3:**
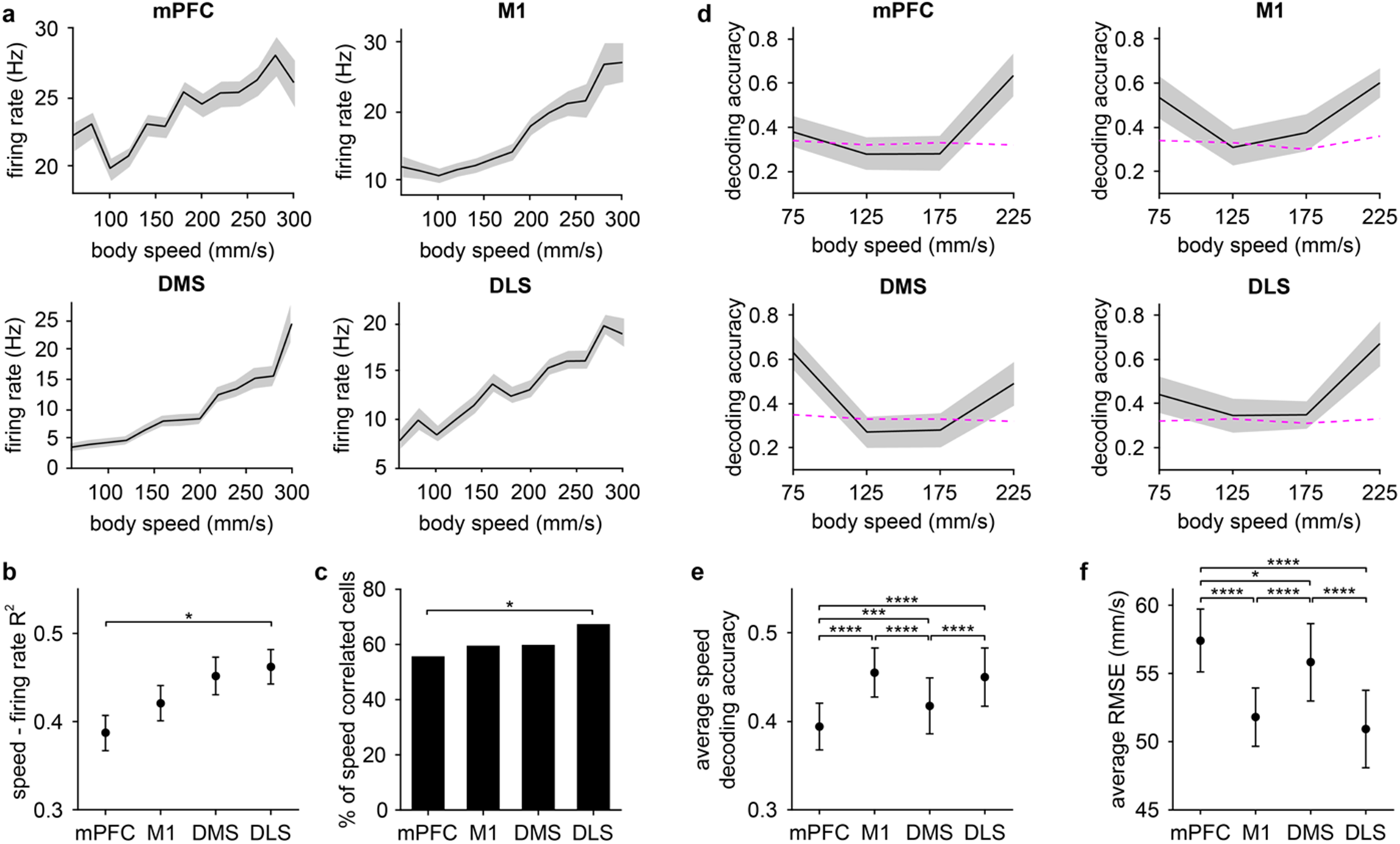
Differential population coding of whole-body speed during walking. **a,** Mean firing rate as a function of body speed of a representative unit from each area. **b,** Mean squared Pearson correlation coefficient (R^2^) between each cell’s firing rate and speed (one-way ANOVA, F_3,853_ = 2.8, p = 0.037). Shading in **a** and **b** represents ± s.e.m. **c,** Percentage of speed correlated cells (n = 122/220 mPFC, 123/207 M1, 129/216 DMS, and 144/214 DLS cells, chi-square test adjusted for 6 comparisons, mPFC vs DLS: p = 0.035). **d,** Mean accuracy of decoding body speed in each area. Magenta dashed line indicates the 95 % confidence interval of decoder performance trained on body speed-shuffled data. **e,** Mean speed decoding accuracy averaged over four speed bins across the four areas (one-way ANOVA, F_3,196_ = 47, p < 0.0001). **f,** Mean root mean squared error (RMSE) of the speed decoder across the four areas (one-way ANOVA, F_3,196_ = 75, p < 0.0001). Data in **d**, **e**, **f** represents the mean ± s.d. of 50 random drawings of 100 cells. Post hoc Tukey-Kramer test was applied for all area-wise comparisons.

### M1 and DLS neurons are more strongly tuned to the gait cycle

Although our analysis so far indicates that the four recorded brain areas differentially represent whole-body speed, it is unclear how these findings relate to the gait cycle at the level of individual limbs—a higher spatial and temporal resolution, providing a more precise measure of the walking process. Previous work has demonstrated that M1 and dorsal striatal neurons display firing rate changes that are tuned to the gait cycle^10, 17^; however, potential regional variations in this phenomenon are unknown. To address this, we used automated pose tracking tools to track the location of limbs as animals behaved in the open field (**Fig. 4a**). During bouts of walking, limbs displayed a characteristic oscillatory motion. Each stride in the gait cycle consisted of a stance phase, when the limb was in contact with the ground, followed by a swing phase, when the limb was raised and propelled forward (**Fig. 4b**). We investigated the relationship between neural activity and the gait cycle. A subset of neurons from all four brain areas exhibited a time-varying change in firing that appeared to be tuned to the rhythmic motion of single limbs during walking (**Fig. 4c**).

**Fig. 4:**
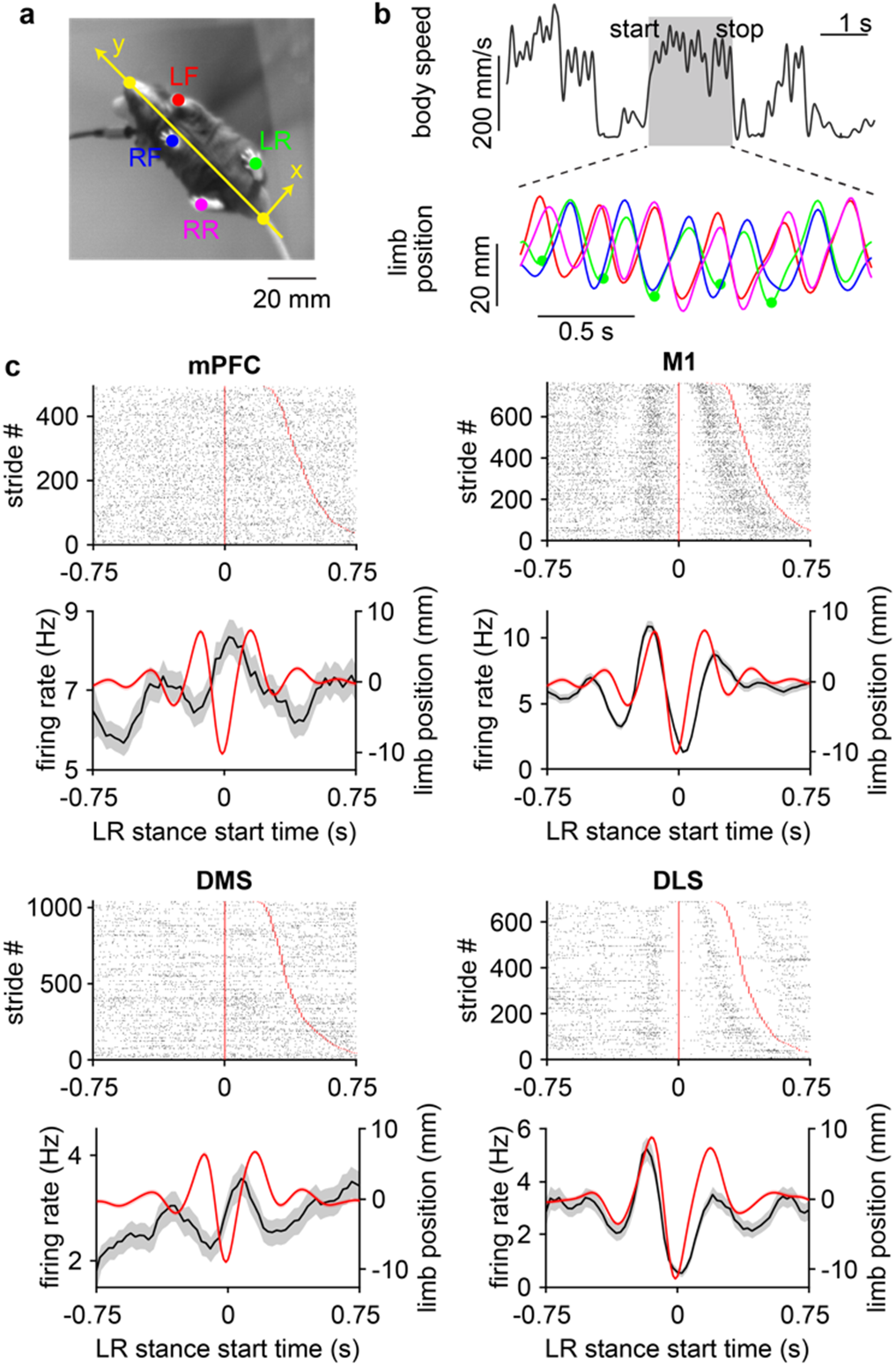
Cortical and striatal neurons are tuned to the gait cycle during walking. **a,** Close-up of a mouse from a video recording displaying the six tracked body features (nose, base of tail, LF: left forelimb, LR: left rear limb, RF: right forelimb, RR: right rear limb). Yellow line represents the direction of forward motion. **b,** Top, whole body speed with a walking bout shaded in gray. Bottom, forward displacement of limb positions during the selected walking bout, colored according to the markers in **a**. The troughs and peaks represent the start of the stance and swing phases of each stride, respectively. Green dots represent the start of LR stance. **c,** Top, spike raster plot of a neuron from each area aligned to LR stance onset. Strides were sorted by increasing duration represented by the red line. Bottom, mean firing rate (black) of the same neurons and mean LR displacement (red) aligned to LR stance onset. Shading represents ± s.e.m.

To characterize the strength of this effect we analyzed spike-limb coupling in the phase domain, by converting limb position to phase (**Fig. 5a**). This transformation enabled us to examine whether spiking activity was preferentially tuned to specific phases of the gait cycle. Neurons from all four areas exhibited limb phase modulated firing patterns (**Fig. 5b**). To quantify the level of modulation we calculated each cell’s spike-phase vector length, a parameter ranging from 0 to 1, indicating the degree of entrainment to the gait cycle of a specific limb^17^. On average, the vector length was similar with respect to all four limbs, though some areas showed a small but statistically significant preference for certain limbs (**Fig. 5c**). Notably, M1 neurons, which were all recorded from the right hemisphere, showed the strongest coupling to left forelimb gait, potentially reflecting a weak contralateral limb preference. To compare spike-limb phase coupling across different brain areas we calculated the limb-averaged vector length. M1 and DLS showed a significantly greater mean vector length than mPFC and DMS (**Fig. 5d**). M1 and DLS also contained a higher proportion of cells with significant phase locking to at least one limb (**Fig. 5e**). These findings indicate that M1 and DLS neurons are not only more strongly tuned to low resolution, whole-body measures of locomotor performance (i.e., speed), but also to rapid, step-by-step limb movements during locomotion.

**Fig. 5:**
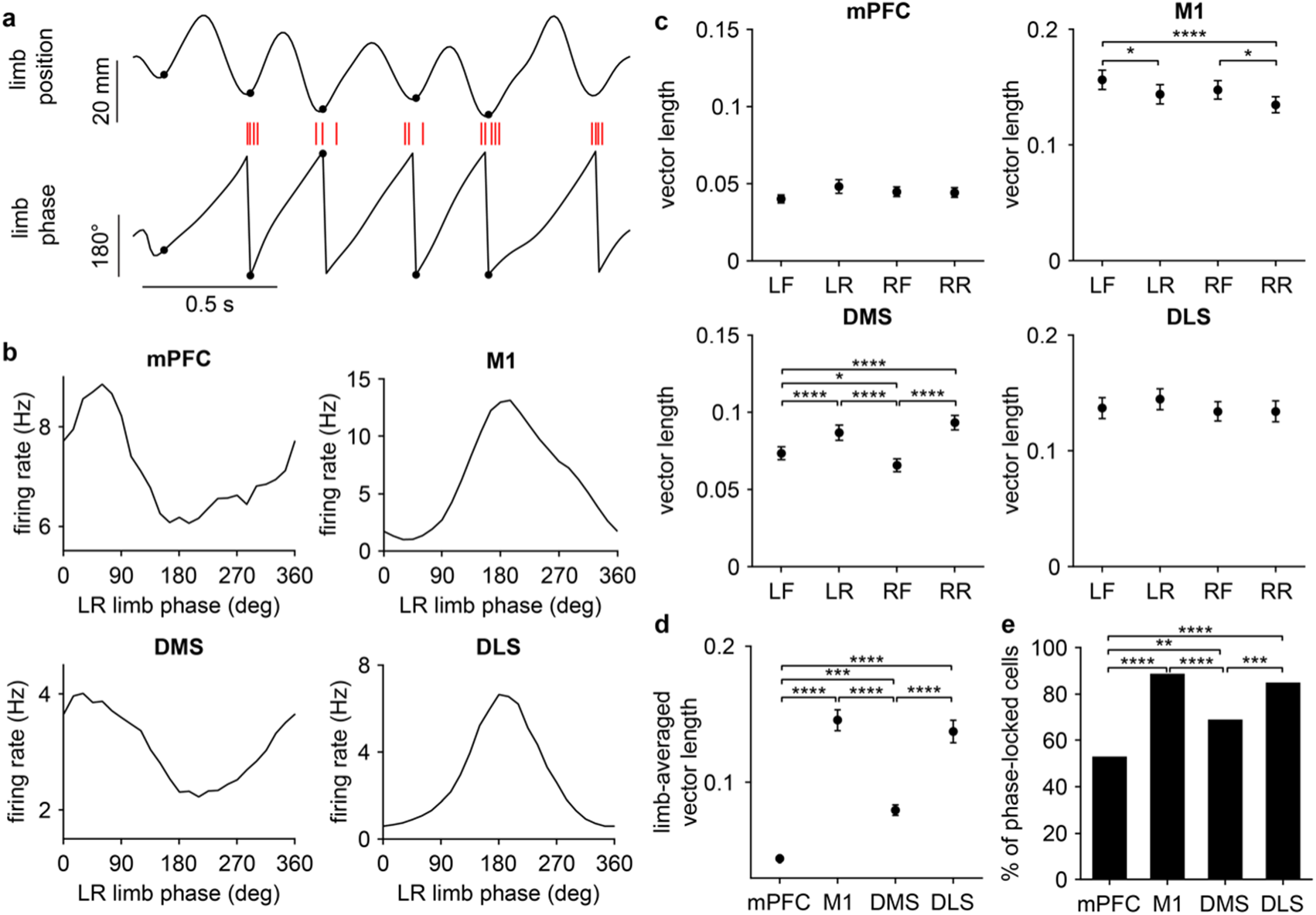
M1 and DLS neurons are more strongly tuned to the gait cycle. **a,** Schematic showing the conversion of limb position to phase. Black markers represent the start of limb stance. Spikes from one neuron are shown as red dashes. **b,** Mean firing rate of a representative neuron from each area aligned to left rear limb (LR) phase. **c,** Mean spike-limb phase vector length compared across each limb (mPFC: n = 220 cells, one-way RM ANOVA, F_3,657_ = 3.2, p = 0.02; M1: n = 207 cells, one-way RM ANOVA, F_3,618_ = 9.6, p < 0.0001; DMS: n = 216 cells, one-way RM ANOVA, F_3,645_ = 29, p < 0.0001; DLS: n = 214 cells, one-way RM ANOVA, F_3,639_ = 1.7, p = 0.18). **d,** Mean limb-averaged spike-limb phase vector length compared across the four areas (one-way ANOVA, F_3,853_ = 64, p < 0.0001). For **c** and **d,** post hoc Tukey-Kramer test was applied for all area-wise comparisons. Data represents mean ± s.e.m. **e,** Percentage of cells in each area with significant phase locking to at least one limb’s gait (n = 117/220 mPFC, 184/207 M1, 149/216 DMS, and 182/214 DLS cells, chi-square test adjusted for 6 comparisons).

### More accurate decoding of gait phase from M1 population dynamics

So far, our results suggest that on average, neurons in M1 and DLS are tuned to single-limb gait at a comparably high level. On the one hand, this aligns with the established role of both areas in motor control as well as the presumed similarity of their dynamics^21, 25^. However, it is unclear if the observed vector lengths, which were calculated across hundreds of trials (i.e., strides), translate to an accurate code for limb phase on a trial-by-trial basis. We therefore next tested whether limb phase could be decoded from neural populations in each of the four brain areas. We trained a decoder to classify neural activity into one of twelve 30 degree phase increments, collectively spanning the entire 360 degree gait cycle (**Fig. 6a**). Decoders trained on M1 or DLS data were both significantly better at predicting the phase of the left forelimb (i.e., contralateral to the recording hemisphere), while mPFC or DMS data showed a weaker limb preference (**Extended Data Fig. 2a**). We confirmed that classification accuracy scaled with population size in all four areas (**Extended Data Fig. 2b**). However, when comparing decoders trained with an equal number of cells, it was evident that the accuracy varied considerably across different regions (**Fig. 6b**). Decoders predicted limb phase below or marginally above chance levels when using mPFC or DMS data, suggesting that population activity in these areas contains a relatively inaccurate representation of gait (**Fig. 6c**). While decoders trained on M1 or DLS data both performed above chance levels, M1 exhibited a significantly more accurate population code (**Fig. 6d**). M1 neural populations also provided the smallest mean absolute error between the predicted and actual phase (**Fig. 6e**). Together, this reveals that among the four recorded areas, M1 contains the most informative population code for gait phase during locomotion.

**Fig. 6:**
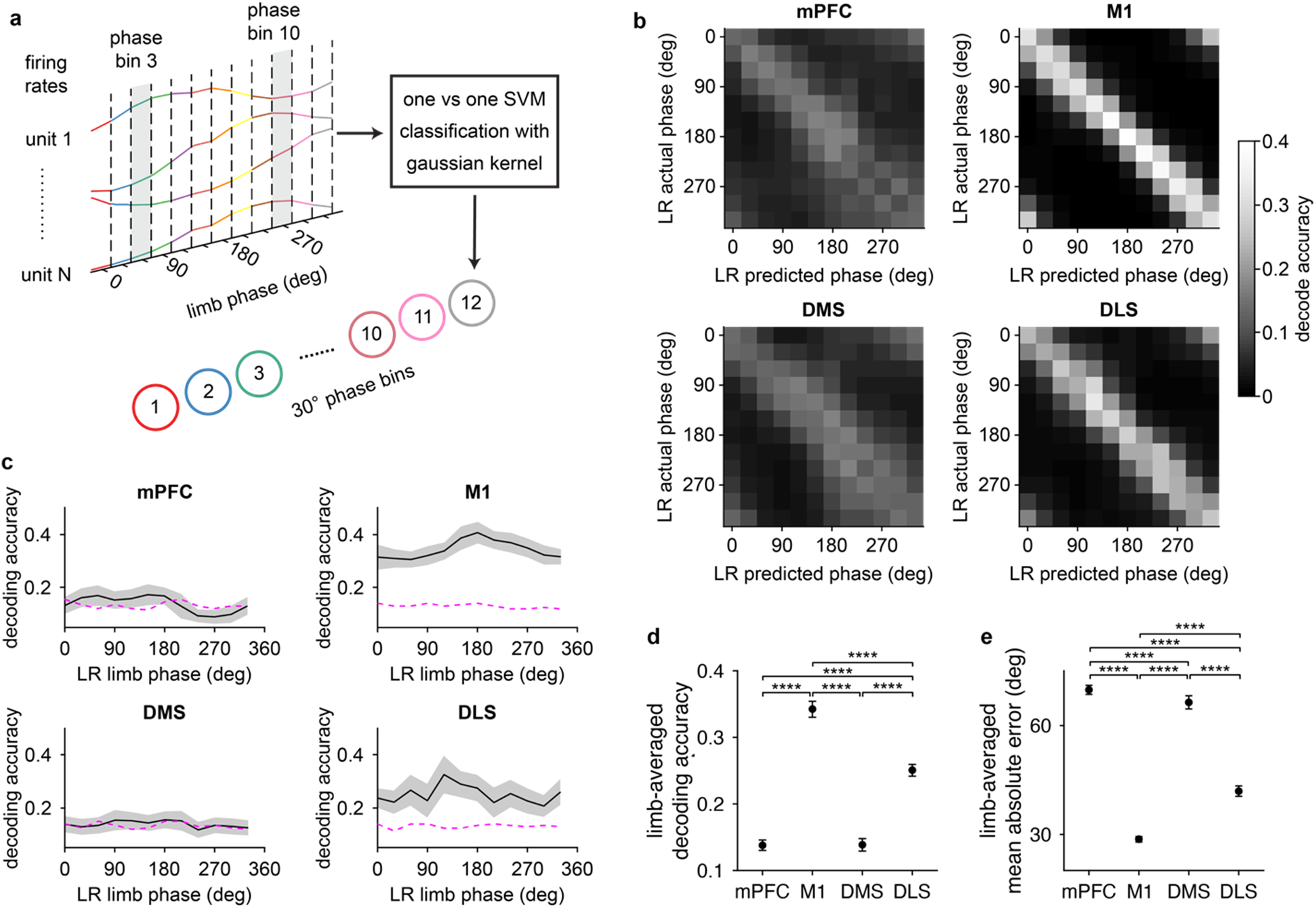
More accurate decoding of gait phase from M1 population dynamics. **a,** Schematic of the approach used to decode the phase of individual limbs undergoing gait using an SVM classifier. Firing rate estimates of single units are binned in 30 degree phase increments for a total of 12 population activity patterns per trial (one trial equals one stride). The decoder is trained to distinguish activity patterns at each phase bin from every other phase bin. **b,** Confusion matrices displaying how well the decoder predicted left rear limb (LR) phase from each area. **c,** Mean decoding accuracy of LR phase. Shaded area represents the s.d. across 50 random drawings of 200 cells. Magenta dashed line indicates the 95 % confidence interval of decoder performance trained on phase-shuffled data. **d,** Mean limb-averaged decoding accuracy compared across the four areas (one-way ANOVA, n = 50 random drawings of 200 cells, F_3,196_ = 5230, p < 0.0001). **e,** Mean limb-averaged absolute angular error of the decoder compared across the four areas (one-way ANOVA, n = 50 random drawings of 200 cells, F_3,196_ = 10551, p < 0.0001). For **d** and **e,** data represents the mean ± s.d. across 50 random drawings of 200 cells. Post hoc Tukey-Kramer test was applied for all area-wise comparisons.

## Discussion

This work challenges the view that striatal circuits simply mirror or contain redundant motor signals from cortex^21, 22^. Instead, the results suggest that the preparatory and performance stages of self-initiated locomotion are preferentially represented by population dynamics in distinct brain areas. These findings have two major implications for understanding the motor functions of cortical and striatal circuits. The first key insight is that despite their strong anatomical connectivity, these areas contain significantly different population codes for locomotion. At a mechanistic level, these differences align with previous work showing that striatal dynamics not only rely on cortical inputs but also on signals integrated from multiple other sources, including local microcircuits, thalamus, and midbrain^43-45^. Second, the results provide evidence for a previously unknown complementary coding scheme, in which DMS population activity more accurately represents the preparatory period, and M1 more accurately represents the performance of locomotion on a step-by-step basis. This regional shift in the encoding of different stages of movement may reflect a functional specialization of these areas, with DMS primarily facilitating the initiation of motion, and M1 primarily regulating limb movements during execution of the gait cycle.

The result that the preparatory time period could be decoded more accurately from DMS neural populations supports previous work implicating the striatum in the initiation and cessation of action sequences^46-48^. Striatal circuits canonically influence movement initiation via the direct and indirect pathways^27, 49^. Both pathways show increased activity in the preparatory phase of motion^14, 50, 51^. Thus, while not directly addressed here, we speculate that both direct and indirect pathway cell populations may reliably encode the preparatory time period, although subtle cell type-specific distinctions have been suggested^17^. In any event, despite an established role for the striatum in movement initiation, it was still surprising to find stronger preparatory signals in striatum relative to cortex. This is because M1 and mPFC are also known to contain robust movement-preparatory neural dynamics and to regulate movement initiation^34-36, 38^. Thus, our work suggests that despite preparatory activity being distributed across cortical and striatal circuits, there are significant regional variations in the quality of this signal as we have seen at the local population level.

But where do these preparatory signals originate? Unsurprisingly, this fundamental question has captivated the field for decades. Single-neuron correlates of movement preparation were first reported in M1, but this was soon followed by similar findings in multiple other areas^6, 52, 53^. As the extent of preparatory activity throughout the brain became apparent, efforts were made to locate the signal’s origins by comparing the latency of individual neuron responses in different areas^8, 12, 34, 54^. However, this approach was inconclusive, yielding only small or inconsistent differences in the timing of preparatory activity. Likewise, our work initially borrowed approaches from these early comparative studies (**Fig. 1f**), but interpreting the proportion of modulated cells was not straightforward. Thus, we substantially built on previous work by turning to population decoding methods to identify the area with the most consistently varying preparatory signal from trial to trial. It is still possible that these signals emerge in cortex before propagating to subcortical circuits including striatum^27^. An alternative mechanism is that preparatory activity may appear earlier in striatal microcircuits and propagate to cortex via basal ganglia-thalamo-cortical feedback loops. There is some supporting evidence that information can flow in both directions depending on task demands, and that signals relevant for learned motor skills flow via feedback loops from striatum to cortex^55^. The origin of preparatory activity in other areas such as cerebellum has also been proposed and cannot be ruled out^54^. Ultimately, this work does not address where this signal originates, but rather where it can be most accurately read out.

Preparatory neural activity occurs in a wide variety of behavioral contexts, but an important consideration in past work has been whether the initial command to move is guided by external factors such as cues or rewards, or generated spontaneously, as in our open field walking task. Indeed, there is evidence that both cortical and striatal dynamics prior to cued movements differ from those of uncued movements^56^. Such differences may occur because cues produce early sensory responses which influence the temporal sequence of signal propagation across the brain^57^. In light of these issues, it is unclear if our findings generalize to other motor functions encountered experimentally. Nevertheless, animals often walk, groom, fidget, and perform other movements spontaneously, requiring an internally generated neural mechanism for initiating such actions. Thus, our work may hold particular relevance for understanding the mechanisms of spontaneous movement initiation, by suggesting a specialized role for striatal dynamics in this process.

After elucidating regional differences in population coding during the preparatory period, we investigated the encoding of locomotor performance. We initially examined speed coding, but this was a relatively low resolution measure of locomotion. Next, tracking individual limbs with high spatial and temporal resolution enabled us to assess how neural activity varied with specific phases of gait. This approach revealed that a sizable fraction of cells in each of the four areas were entrained to the gait cycle. We again found significant regional differences in the ability to decode gait phase from neural populations. However, rather than pointing to DMS, these results indicate that M1 contains the most accurate population code for gait. Taken together, this work demonstrates that locomotor preparation and performance are more accurately encoded in striatal and cortical circuits, respectively.

Functionally, the coupling of M1 activity to the gait cycle may signify a motor output signal, coordinating with brain stem and spinal cord central pattern generator circuits to fine-tune locomotion^9, 58^. However, unlike humans, where M1 damage severely impair locomotion^59^, M1 lesions in rodents appear to transiently disrupt fine limb movements (e.g., reaching and manipulating small objects or climbing over obstacles) while not appreciably impairing walking on a flat surface^60-63^. This suggests that rather than directly controlling gait, in rodents, this phasic activity may represent a form of efference copy or sensory feedback, used for detecting and correcting deviations from predicted locomotor rhythms. Yet until now the relative extent of gait-modulated dynamics in different cortical and striatal regions was not well characterized. Our work revealed a marked contrast in the strength of gait-related information coding between M1 and mPFC, as well as between DLS and DMS, suggesting this information is enhanced in areas associated with locomotor performance. In terms of whether these findings generalize to other motor functions, a recent study found that population decoding of continuous forelimb kinematics in a reach/pull task was significantly better in mouse M1 relative to mPFC, again supporting a stronger role for M1 in performing limb movements^22^. However, that study also found similar decoding accuracy between M1 and dorsal striatum. If this discrepancy is due to distinct behavioral task demands, this would again suggest a need to carefully dissociate self-initiated from instructed movement mechanisms^56, 64^. While DMS and M1 respectively showed the most accurate population code for locomotor preparation and performance, it is noteworthy that another striatal subregion, DLS, consistently appeared as the area with the second most accurate code. This may signify a specialized role for this area in integrating a variety of behaviorally relevant signals. Along these lines, previous work has shown that DLS “multiplexes” several different types of kinematic information^65^.

Lastly, what is the potential advantage of a complementary neural coding scheme, in which distinct areas more accurately encode locomotor preparation and gait? Fundamentally, the regional differences in decoding quality may reflect a refinement of movement-related temporal dynamics as they propagate across different brain areas^32, 40^. At the same time, our results suggest that signals for locomotor preparation and performance may be refined or amplified in distinct circuits. Such a complementary coding scheme may offer an efficient strategy for minimizing crosstalk between neural signals for motor preparation and those for the continuous production of movements. An alternative strategy proposes that the same neural populations toggle between computing preparatory and performance-related information^66^. It is also intriguing to consider the possibility that the coding differences we identified reflect functional differences as well, with DMS specialized for preparing to initiate, and M1 specialized for performing locomotion. However, while sometimes implicitly assumed, it is not yet known whether a more accurate information code necessarily translates to a stronger behavioral role for that population. In spite of this ambiguity, comparing neural codes may help to clarify and contextualize the behavioral contribution of different brain circuits—especially those with closely related functions.

**Extended Data Fig. 1:**
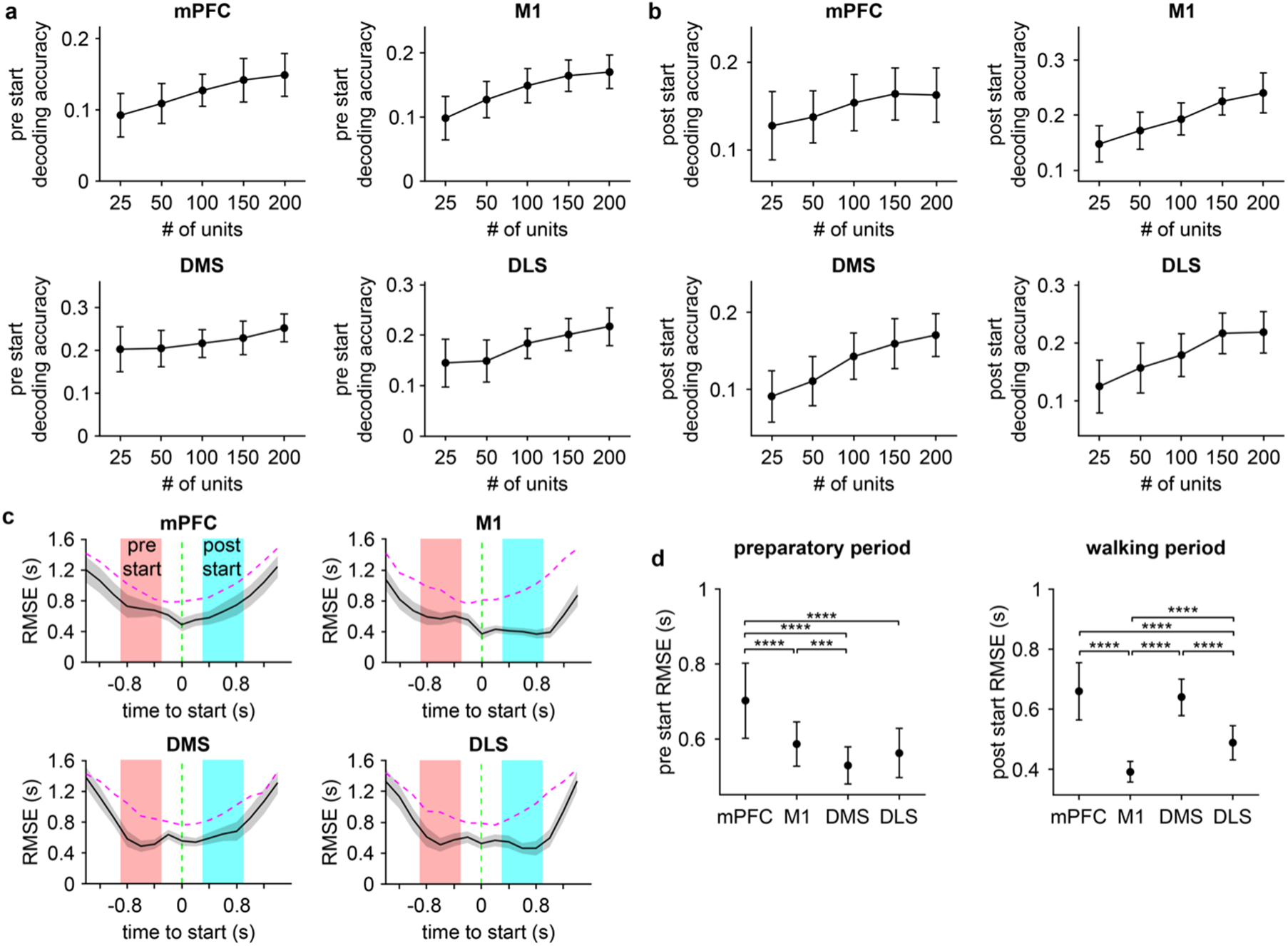
Population decoding of time in the preparatory and performance periods of locomotion. **a,** Mean decoding accuracy in the preparatory (“pre-start”) period as a function of the number of neurons. mPFC: one-way ANOVA, F_4,245_ = 33, p < 0.0001; M1: one-way ANOVA, F_4,245_ = 54, p < 0.0001; DMS: one-way ANOVA, F_4,245_ = 13, p < 0.0001; DLS: one-way ANOVA, F_4,245_ = 35, p < 0.0001. **b,** Mean decoding accuracy in the performance (“post-start”) period of walking as a function of the number of neurons. mPFC: one-way ANOVA, F_4,245_ = 12, p < 0.0001; M1: one-way ANOVA, F_4,245_ = 71, p < 0.0001; DMS: one-way ANOVA, F_4,245_ = 57, p < 0.0001; DLS: one-way ANOVA, F_4,245_ = 51, p < 0.0001. For **a** and **b**, data represents the mean ± s.d. across 50 random drawings of the specified number of cells. **c,** Mean decoder root mean squared error (RMSE) aligned to the start of walking. Shaded area represents the s.d. across 50 random drawings of 200 cells. Magenta dashed line indicates the 5 % confidence interval of RMSE for time-shuffled data. Green dashed line represents the onset of walking. **d,** Left, mean RMSE in the preparatory period compared across the four areas (one-way ANOVA, n = 50 random drawings of 200 cells, F_3,196_ = 56, p < 0.0001). Right, mean RMSE in the performance period of walking compared across the four areas (one-way ANOVA, n = 50 random drawings of 200 cells, F_3,196_ = 191, p < 0.0001). Data represents the mean ± s.d. across 50 random drawings of 200 cells. Post hoc Tukey-Kramer test was applied for all area-wise comparisons.

**Extended Data Fig. 2:**
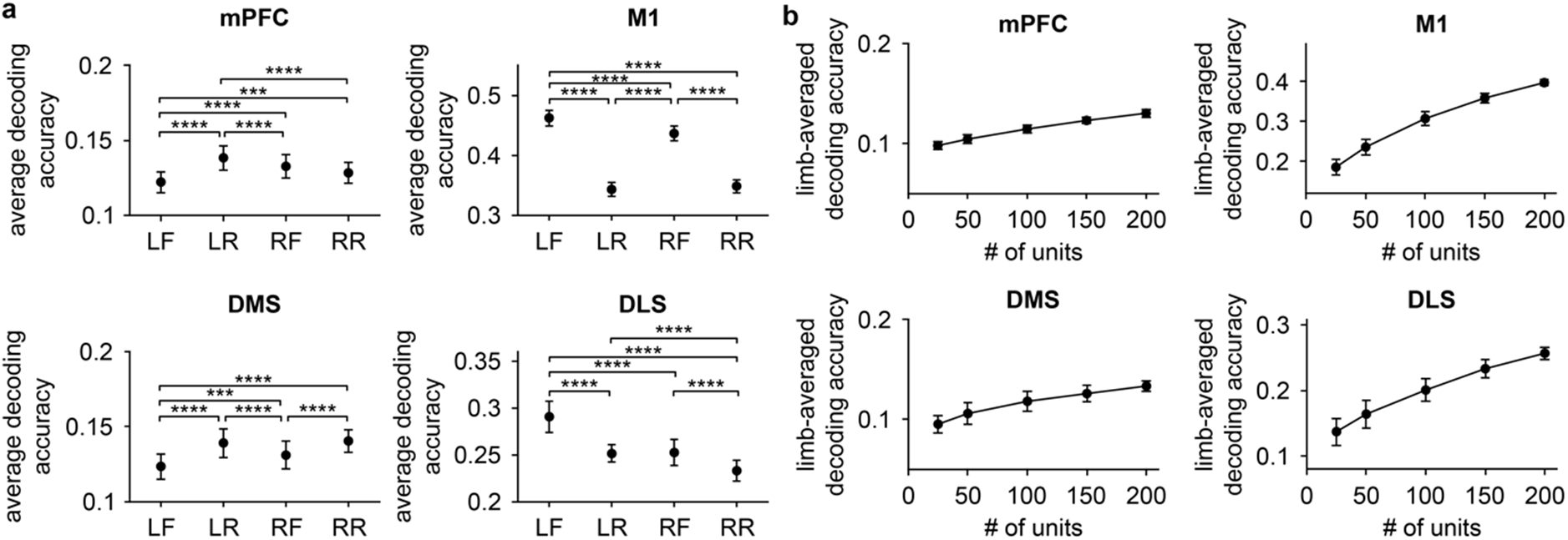
Population decoding of gait phase. **a,** Mean decoding accuracy averaged over the gait cycle compared across the four limbs. mPFC: one-way RM ANOVA, F_3,147_ = 49, p < 0.0001; M1: one-way RM ANOVA, F_3,147_ = 1397, p < 0.0001; DMS: one-way RM ANOVA, F_3,147_ = 44, p < 0.0001; DLS: one-way RM ANOVA, F_3,147_ = 271, p < 0.0001. Data represents the mean ± s.d. across 50 random drawings of 200 cells. Post hoc Tukey-Kramer test was applied for all limb-wise comparisons. **b,** Mean limb-averaged decoding accuracy as a function of the number of neurons. mPFC: one-way ANOVA, F_4,245_ = 589, p < 0.0001; M1: one-way ANOVA, F_4,245_ = 1453, p < 0.0001; DMS: one-way ANOVA, F_4,245_ = 152, p < 0.0001; DLS: one-way ANOVA, F_4,245_ = 410, p < 0.0001. Data represents the mean ± s.d. across 50 random drawings of the specified number of cells.

## Methods

### Animals

All procedures were approved by the University of California, Los Angeles Chancellor’s Animal Research Committee. Single housed male C57BL/6J mice, 11-24 weeks old (The Jackson Laboratory) at the time of the surgery were used in the experiments. Animals were kept on a 12 hour light cycle, and group housed until the day of the surgery.

### Surgical procedures

Animals underwent a surgical procedure under aseptic conditions and isoflurane anesthesia on a stereotaxic apparatus. The procedure involved attaching a custom 3D printed head cap on the skull, and drilling a craniotomy window. We then implanted a single-shank 64 electrode silicon microprobe^67^ (model 64D-sharp, Masmanidis lab) which was superglued to a microdrive shuttle (Nano Drive, Cambridge Neurotech). The probes were electrically wire bonded to a flexible cable that was connected to a miniature electrophysiological head stage (HS-64m, White Matter LLC). The head cap contained a base plate which was securely fixed to the skull using dental cement, and a cap which was attached to the base plate with a pair of screws. The head cap featured space for the flexible cable, connected to the head stage. The silicon probe was implanted in the right hemisphere targeting one of three brain regions: M1 (n = 7 mice, probe tip at 0.74 mm anterior, 1.4 mm lateral, 1.05 mm ventral relative to bregma), DLS (n = 4 mice, probe tip at 0.74 mm anterior, 2.2 mm lateral, 2.1 mm ventral relative to bregma), and mPFC (n = 2 mice, probe tip at 1.6 mm anterior, 0.3-0.45 mm lateral, 1.05 mm ventral relative to bregma). Since the probe was movable using the microdrive, in some mice, after completing M1 recordings we moved the probe down to record from DMS (n = 4 mice, probe tip at 0.74 mm anterior, 1.4 mm lateral, 2.05 mm ventral relative to bregma). After inserting the silicon probe, a layer of Vaseline was applied around the exposed silicon probe. A layer of UV curable cement was then applied to fix the microdrive to the skull. A craniotomy was made over the left hemisphere for the insertion of a silver ground wire.

### Motion tracking

Behavior was monitored in a 60 cm x 60 cm open field arena, featuring a transparent floor for bottom-up imaging using a high-speed camera (acA2040-90umNIR, Basler). The arena was housed in a darkened room and illuminated by infrared light. Video was captured at 80 fps and a spatial resolution of 0.3 mm per pixel using Streampix 8 software (Norpix). To synchronize video with electrophysiological data, the camera was triggered by a shared 80 Hz clock signal generated from a data acquisition card (SB-6356, National Instruments). Prior to the first recording session, animals underwent a habituation phase in which they were allowed to freely explore the arena for 15 minutes per day over two to three days. Each recording session was 30 minutes in duration. The arena was cleaned between recording sessions by wiping with distilled water and ethanol. Mice typically displayed a variety of spontaneous behaviors including resting, grooming, and walking. Limb movements were tracked offline using SLEAP software^68^, which was trained to detect the 2D coordinates of the four limbs, nose, and base of the tail. Whole-body speed was calculated as the root mean squared value of the four individual limb speeds.

We performed a semi-automated identification of walking bouts in order to eliminate periods when animal movement did not correspond to locomotion (e.g., grooming). This step involved replaying the tracked body coordinates during candidate walking frames (initially identified automatically by applying a speed threshold), and manually accepting only frames in which the limbs displayed a clear rhythmic motion characteristic of gait. After obtaining a preliminary estimate of the onset time of each walking bout, the onset time was further refined by identifying the peak in body acceleration within a 50 frame (0.625 s) window centered around the estimated onset time. If no peak acceleration with a prominence above 1000 mm/s^2^ was found in this window, the walking bout was discarded. We further removed walking bouts that were less than 1 s apart. The minimum duration of usable walking bouts was 1 s, with shorter bouts being discarded. Spatial coordinates were then smoothed with a third order Savitzky-Golay filter. To identify the onset time of individual strides, limb positions were projected onto the nose-tail axis, and bandpass filtered from 0.5 to 8 Hz. This filtered signal revealed the cyclical motion of limbs during gait. Each stride begins with the stance phase, whose onset corresponds to the minima in the gait cycle (indicating that the limb is closest to the nose). The stance is followed by the swing phase, whose onset corresponds to the maxima in the gait cycle (indicating that the limb is furthest from the nose). The end of the swing coincides with the start of the next stride’s stance. The limb phase angle was calculated by applying a Hilbert transform on the filtered limb-to-nose distance signal.

### Electrophysiology

Before each recording session, the head stage in the head cap was connected to a flexible wire tether via an in-line electrical rotary joint (White Matter LLC), allowing the animal to move freely in the arena. Electrophysiological data were sampled at 25 kHz per channel. The microdrive was used to lower the silicon probe by a depth of 0.0625 to 0.5 mm between sessions. Between 9 and 21 recording sessions were carried out in each of the four brain areas of interest. Offline, raw data were bandpass filtered from 600 to 7000 Hz, spike sorted with Kilosort^69^ and manually curated with Phy.

### Identification of units modulated around the onset of walking

For each unit, spikes within a ±1 s window around the onset of walking were binned in 25 ms increments. The resulting spike counts for each trial were convolved with a Gaussian kernel (s.d. = 0.1 s), and then averaged into 0.1 s bins. We used a paired t-test to compare firing rates at each time bin with the rate in the baseline period, defined as -1 to -0.9 s from movement onset. We then evaluated the percentage of units with statistically significant (p < 0.05) deviations from baseline firing at each time bin.

### Population decoding of time around the onset of walking

Walking bouts as identified earlier were used as the trials for decoding. For each trial, neural population activity was examined within a ±1.5 s window around walking onset. Single unit firing rates were calculated from spike counts in 25 ms increments and convolved with a Gaussian kernel (s.d. = 0.1 s), and then averaged into 0.2 s bins. We constructed 15 population firing rate vectors per trial, with each vector corresponding to a non-overlapping 0.2 s time bin within the ±1.5 s window. To pool data across sessions and animals, we randomly selected 50 trials from all sessions. Time was decoded from the population firing rates in each trial by using a decoder tasked with classifying each rate vector in the trial to one of the 15 distinct time bins^40^. This classification task employed a multiclass SVM with a Gaussian kernel, implemented via the LIBSVM library in MATLAB. The SVM applied a one-against-one multiclass approach, training binary classifiers to distinguish firing patterns between all pairs of time bins, yielding 105 classifiers in total. Each classifier produced a score reflecting how closely a test vector matched the neural activity pattern of a specific time bin. The time bin with the highest score was selected as the predicted time point. Decoder performance was optimized for each brain region by tuning the SVM’s regularization parameters (Box Constraint and Kernel Scale) through grid search with fivefold cross-validation, capped at 20 evaluations.

To evaluate decoder performance, we conducted 50 decoding iterations, each leaving one random trial out for testing, using a set of N neurons. To generate an accuracy distribution for statistical analysis, 50 random draws of N units were performed, with N set to 200 units for the primary analysis. The effect of population size on decoding accuracy was assessed by training models on random samples of 25, 50, 100, 150, and 200 neurons from each brain region. To generate shuffled control data, we performed the same decoding analysis on firing rates that were randomly reassigned to different time bins within the same trial. Decoder accuracy distributions from time bin-shuffled models were used to calculate 95 % confidence intervals. RMSE was computed by comparing actual and predicted time bins. For assessing performance relative to walking onset, decoding accuracy and RMSE were averaged over two intervals: pre-start (0.3 to 0.9 s pre onset) and post-start (0.3 to 0.9 s post onset).

### Identification of speed correlated units

Speed correlated units were identified based on whole-body speed data during bouts of walking. Speed data were binned in increments of 20 mm/s from 50 to 310 mm/s, and for each cell the average firing rate was calculated at each speed increment. The speed correlation score represented the square of the Pearson correlation coefficient (R^2^) between each cell’s firing rate and speed. Cells with a statistically significant correlation (p < 0.05) were classified as speed correlated.

### Population decoding of speed

In this analysis, neural population activity binned into body speeds was used as trials for decoding. Neural population activity was analyzed for each trial across four ranges of body speed during walking: 50 – 100 mm/s, 100 – 150 mm/s, 150 – 200 mm/s, and 200 – 250 mm/s. Single unit firing rates were calculated from spike counts in 25 ms increments and convolved with a Gaussian kernel (s.d. = 0.1 s), then averaged into 0.2 s bins. Concurrently, average body speed was calculated for the same time bins. To combine data from multiple sessions and animals, we randomly selected 100 trials per speed bin from all sessions. A similar SVM decoder used for predicting time around walking onset was applied, but here it was configured for four output classes representing body speeds. Decoder performance was evaluated by leaving out two random trials in each of 50 models trained on 100 selected neurons. To build an accuracy distribution for statistical analysis, 50 random samples of 100 neurons were drawn. For control, shuffled data were generated by randomly assigning each neuron’s firing rate estimate to a different speed bin. RMSE was calculated by comparing predicted body speeds with actual speeds. Decoder accuracy distributions from speed bin-shuffled models were used to calculate 95 % confidence intervals.

### Identification of gait phase-locked units

For all strides, single unit spikes were counted at each phase from 0 to 360° in 15° increments. This resulted in each single unit having 24 rates across the full 360° gait cycle. The mean vector length was calculated from the normalized average of these 24 rates using circular statistics^17^. The mean vector length, ranging from 0 to 1, reflects the degree of neural entrainment to the gait cycle. We performed a spike time jitter analysis by shifting spike times by a random amount within ±0.15 s and recalculating the mean vector length over 1000 iterations. Neurons were classified as phase-locked to gait if their actual vector length exceeded 95 % of the spike jittered vector lengths for at least one limb’s phase.

### Population decoding of gait phase

Neural population activity was analyzed for each trial (i.e., stride) across twelve 30° phase bins spanning the full 360° gait cycle. Single unit firing rates were calculated from spike counts in 25 ms increments and convolved with a Gaussian kernel (s.d. = 0.1 s), then averaged into 0.2 s bins. To combine data from multiple sessions and animals, we randomly selected 200 trials per phase bin from all the sessions. A similar SVM decoder used for predicting time around walking onset was applied, but here it was configured with 12 output classes representing single limb phase bins. Separate models were trained to predict the gait phase for each of the four limbs. Decoder performance was evaluated by testing on four randomly withheld trials in each of the 50 models, each trained on a set of N units. To generate an accuracy distribution for statistical analysis, 50 random draws of N units were performed, with N set to 200 units for the primary analysis. To investigate how population size influenced model performance, additional models were trained using random subsets of 25, 50, 100, and 150 units from each area. For control, each neuron’s firing rate estimate in a given phase bin was replaced with its firing rate estimate from a randomly chosen phase bin. Decoder accuracy distributions from phase bin-shuffled models were used to calculate 95 % confidence intervals. The mean absolute error was defined as the absolute angular difference between actual and predicted limb phases.

### Quantification and statistical analysis

Neural data were pooled across all recording sessions in each brain area. Statistical analysis was performed using MATLAB code. The sample size, type of statistical test and P values are indicated in the figure legends. Data distribution was assumed to be normal, but not explicitly tested. The t-tests were two-sided. One-way ordinary or repeated measures analysis of variance (ANOVA) was followed by Tukey’s post hoc test for multiple comparisons. Significant differences throughout all the figures are represented by *p<0.05, **p<0.01, ***p<0.001, ****p<0.0001.

## Data availability

The data that support the findings of this study are available at: https://doi.org/10.5281/zenodo.14969115

## Code availability

The code used for analyzing data is available at: https://github.com/sotmasman/Complementary-ctx-str-encoding

## Acknowledgments

This work was supported by an American Heart Association predoctoral fellowship 25PRE1357956 to D.S., NIH grants R01NS136137, R01NS136123 and R01NS125877 to S.C.M., and NSF CAREER 1943467, NIH grants DP2NS122037 and R01NS121097 to J.C.K.

